# Investigating The Potential of Division of Labour in Synthetic Bacterial Communities for the Production of Violacein

**DOI:** 10.1101/2025.01.07.631562

**Authors:** Harman Mehta, Jose Jimenez, Rodrigo Ledesma-Amaro, Guy-Bart Stan

## Abstract

With advancements in synthetic biology and metabolic engineering, microorganisms can now be engineered to perform increasingly complex functions, which may be limited by the resources available in individual cells. Division of labour in synthetic microbial communities offers a promising approach to enhance metabolic efficiency and resilience in bioproduction. By distributing complex metabolic pathways across multiple subpopulations, the resource competition and metabolic burden imposed on an individual cell is reduced, potentially enabling more efficient production of target compounds. Violacein is a high-value pigment with anti-tumour properties that exemplifies such a challenge due to its complex bioproduction pathway, imposing a significant metabolic burden on host cells. In this study we investigated the benefits of division of labour for violacein production by splitting the violacein bioproduction pathway between two subpopulations of *Escherichia coli* based synthetic communities. We tested several pathway splitting strategies and reported that splitting the pathway into two subpopulations expressing VioABE and VioDC at a final composition of 60:40 yields a 2.5 fold increase in violacein production as compared to a monoculture. We demonstrated that the coculture outperforms the monoculture when both subpopulations exhibit similar metabolic burden levels, resulting in comparable growth rates, and when both subpopulations are present in sufficiently high proportions.

## Introduction

Synthetic biology aims to engineer existing and develop new biological systems that can perform novel and useful functions. Recent progress in the field has vastly enhanced our capability of engineering organisms to perform desired functions. Historically, synthetic biology and biotechnology predominantly focus on modifying single organisms to carry out desired functions in a consolidated bioprocess. However, engineering additional functionality into a living cell can increase metabolic burden, the burden on the cell due to competition for cellular resources and energy, which leads to reduced growth rates, reduced expression rates and provokes genetic instability making engineered cells more prone to negative selection than wild type strains [15], [14]. In nature, cells seldom grow in isolation, and rather exist in diverse interacting communities [12]. This has been shown to confer many advantageous properties to the members of these consortia. Microbial communities demonstrate increased robustness to environmental perturbations and resilience to mutation [2]. There is extensive exchange of metabolites between the members of the communities, with reduced metabolic burden on individual members and increased cooperation [2], [12]. Inspired by these properties of natural microbial consortia, synthetic microbial cocultures are being developed to overcome the limitations of monocultures [18], [20],[19]. Synthetic microbial cocultures can overcome the limitations of metabolic burden in engineered monocultures by performing division of labour, where complex pathways are distributed between multiple cells. Synthetic cocultures implementing division of labour also reduce the burden due to heterologous enzyme expression [18]. Division of labour can also allow modulation of sections of the pathway, by controlling the activation of different reaction, expressing them in specialist organisms at desired growth rates and subpopulation ratios [13], [17].

Violacein is a purple-hued pigment produced by a wide range of naturally occurring bacteria found in a variety of environments ranging from deep seas, to forests and even polar glacial reserves [21], [8], [4]. It has been demonstrated to possess anti-cancer [10] and anti-bacterial properties particularly against gram positive bacteria [21], [7] and is also used to prevent and treat stomach ulcers [21], [7]. Due to its strong colouration, it is also used as a bio-dye [1]. It has also been shown to have antimicrobial properties against some antibiotic resistant bacterial strains of *Staphylococcus aureus* [7]. Due to its extensive applications but limited yields in bioproduction, it is a high value product. Violacein is produced from tryptophan by a five step enzymatic pathway through the expression of the genes *vioABCDE* (Figure 1A). The production of violacein in *E. coli* has been demonstrated to have significant growth deficits on the cell, showing that the heterologous gene expression of the pathway is burdensome on the cell [6].

**Figure 1:**
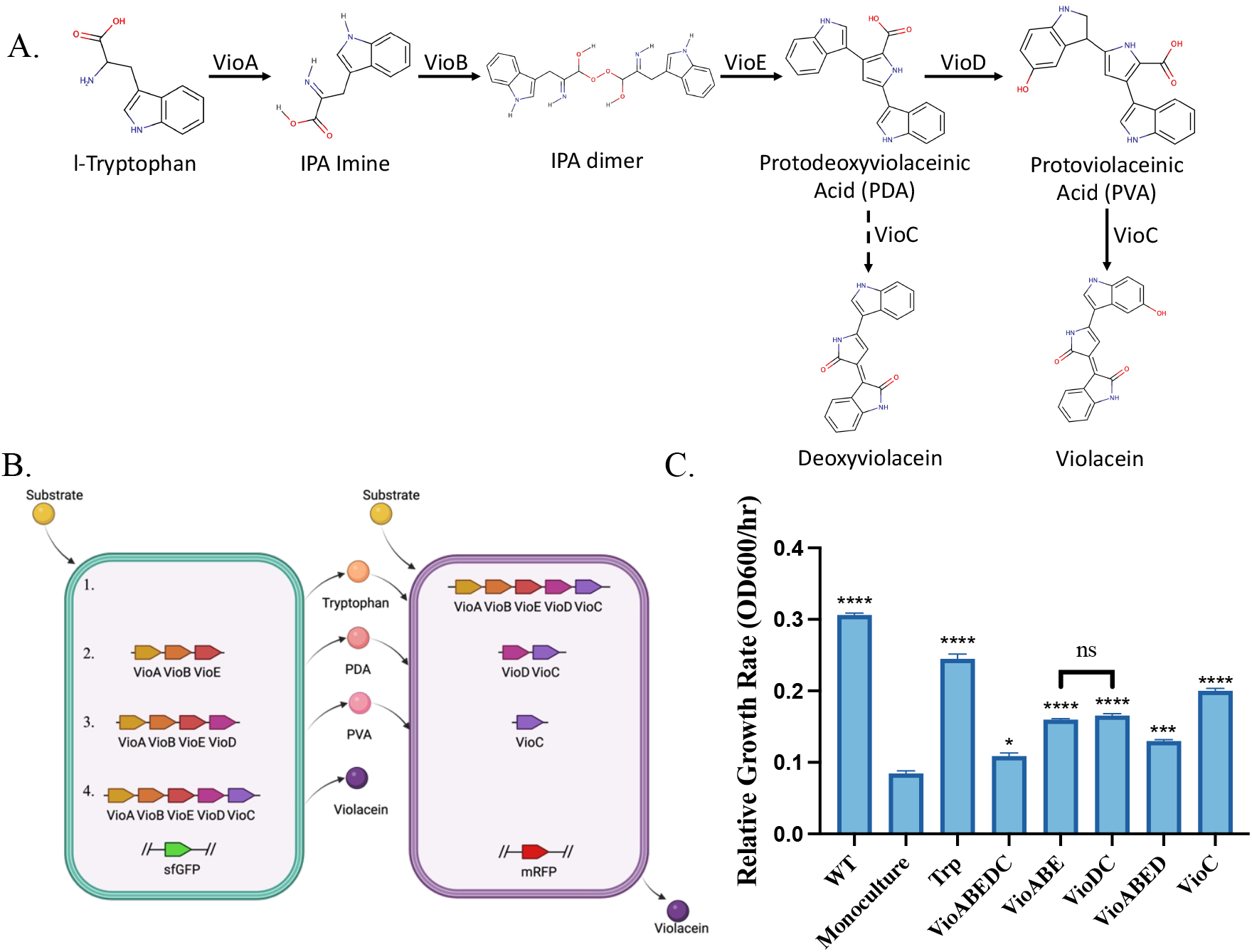
A. Violacein bioproduction pathway. The five gene pathway converts tryptophan to violacein and deoxyviolacein. Abbreviation: IPA: 2-Imino-3-(indol-3-yl)propanoate [3]. B. Violacein pathway split strategies. The proposed pathway splitting strategies for violacein production are demonstrated in the two subpopulations. The green cell represents a tryptophan overproducing strain used to produce violacein precursors. Strategy 1 refers to the pathway splitting at tryptophan. Strategy 2 refers to the pathway splitting at PDA. Strategy 3 refers to the pathway splitting at PVA. Strategy 4 shown here is the production of violacein using a monoculture, to be used as a control for coculture experiments. C. Growth rates of the strains developed in this study, calculated mid log phase in a microplate reader. The monoculture has significant growth defects, compared to the wild type strain. The VioABE and VioDC strains have similar growth rates, and show significant improvement in the growth rates compared to the monoculture. Relative growth rates are calculated at mid log phase of growth (OD600). Values shown are n=3 biological replicates with mean value shown and error bars representing standard deviation. Statistically significant differences were determined using two-tailed Student’s t-test (* represents p *<* 0.05, ** represents p *<* 0.01, *** represents p *<* 0.001, **** represents p *<* 0.0001, ns represents not significant). Symbols above the bars represent the statistical significance of the difference with the monoculture. OD600 measured using a microplate reader.

In this study, we investigated the advantages of division of labour in the context of the violacein biosynthetic pathway expressed in an *E. coli* based two member community. We investigated different pathway splitting points and different coculture compositions and identify conditions where the coculture performs better than the monoculture.

## Materials and Methods

### Bacterial strains and plasmids

*E. coli* strains DH10B (F-*mcrA* Δ(*mrr-hsdRMS-mcrBC*) *ϕ*80*lacZ* ΔM15 Δ*lacX74 recA1 endA1 araD139* Δ(*ara-leu*)7697 *galU galK λ*– *rpsL(StrR) nupG*) and DH5*α* (F – *ϕ80lacZDM15* Δ(*lacZYA-argF*) *U169 recA1 endA1 hsdR17* (rk–, mk+) *phoA supE44* l – thi–1 *gyrA96 relA1*) were used for cloning of all violacein constructs and was grown in Luria Broth (LB) media at 37^°^C with appropriate antibiotics (100 *µ*g/mL Ampicillin, 10 *µ*g/mL Tetracycline, 50 *µ*g/mL Kanamycin). For expression of the violacein plasmids, BL21 (DE3) (F – *ompT hsdSB* (rB-mB-) *gal dcm* (DE3)) was used. For tryptophan overproduction, *E. coli* strain, B6, BL21 (DE3) (F – *ompT hsdSB* (rB-mB-) *gal dcm* (DE3) Δ*trpR* Δ*tnaA*) was received from Prof. Zhang at Tsinghua University [9]. The violacein genes were obtained from the Koffas Lab [11] (pETM6-G6-vioABECD)on (Addgene)(Addgene plasmid # 66536 ; http://n2t.net/addgene:66536 ; RRID:Addgene 66536). The two populations for cocultures were tagged with sfGFP and mRFP1 expressed constitutively in plasmids. The plasmids created in this study are detailed in Supplementary Table 1. The bacterial strains used in this work are detailed in Supplementary Table 2. Polymerase chain reactions, Gibson and Golden Gate assemblies were used to build those plasmids. All plasmid sequences were verified using Sanger sequencing and Oxford Nanopore whole plasmid sequencing. The plasmid for tryptophan over production, pHM068, the TrpED genes were amplified from the *E. coli* MG1655 genome using the oligos TrpED Fwd and TrpED Rev. Two point mutations were introduced using around-the-world PCR with the oligo pairs Ser40LeuT Fwd - Ser40LeuT Rev, and Met293ThrC Fwd - Met293ThrC Rev. The plasmid was constructed with the TrpED cassette and a sfGFP cassette using Start-Stop Golden Gate assembly [16]. The violacein genes were amplified along with the low strength T7 mutant promoter, G6, and RBS from the pETM6-G6-vioABECD plasmid using the oligos mentioned in Supplementary Table 3. The plasmids were constructed using restriction digestion and ligation. The plasmid pHM120 was constructed with mRFP using Start-Stop Golden Gate Assembly [16].

### Culture conditions

All experiments described in this article were conducted in Lysogeny Broth, with appropriate antibiotics (100 *µ*g/mL Ampicillin, 10 *µ*g/mL Tetracycline, 50 *µ*g/mL Kanamycin). The experiments were conducted at two different volumes, 5mL of liquid culture in 14 mL round bottom culture tubes and 10 mL of liquid culture in 50 mL conical flasks. From overnight liquid cultures, fresh subcultures were made at OD600 of 0.01 and inoculated at 37^°^C for 5 hours. After 5 hours, 0.1 M Isopropyl *β*-d-1-thiogalactopyranoside (IPTG) was added to the cultures and for the 5 mL cultures in 14 mL tubes, the lids were replaced by autoclaved foam stoppers for consistent and sufficient aeration. The cultures were then incubated at 20^°^C for 19 hours.

The growth experiments were conducted in clear flat-bottomed 96 well plates in a TECAN Spark microplate reader. From overnight liquid cultures of the strains, fresh subcultures were made in M9 media supplemented with 10% casamino acids and 0.8% glucose. To each well 200 *µ*L of culture volumes were added with starting OD600 of 0.025 and the plate was covered with a Breath-Easy membrane (Diversified Biotech). The plates were grown in the plate reader for 16 hours with readings every 15 minutes at 37^°^C with double orbital shaking, 1.5 mm amplitude. For each strain, three biological replicates were tested and growth curves were fitted to a Gompertz curve, which was used to calculate the growth rate at mid log phase.

### Metabolite purification and quantification

For the quantification of violacein and deoxyviolacein, at the end of the experiment after the cultures were incubated at 20^°^C for 19 hours, the metabolites were extracted in absolute methanol. 1 mL of liquid culture was pelleted in a 1.5 mL microcentrifuge tube in a bench-top centrifuge and the supernatant was discarded. The pellet was resuspended in 1 mL absolute methanol and transferred to a 2 mL screw cap microtube with glass beads. The tubes were shaken at 6000 rpm for 5 minutes, with 15 second breaks every minute, using a Precellys Evolution Homogenizer. The samples were spun down in a bench-top centrifuge and the supernatant was transferred to a HPLC vial. The samples were analysed in a Vanquish Core HPLC system with the YMC Carotenoid C30 column (150 × 4.6 mmI.D. S-3 *µ*m). The mobile phases used were acetonitrile (A) and water (B), both containing 0.1% formic acid. The mobile phase was run with a flow rate of 1 mL/min with the following gradient: 0 min, 5% A; 1 min, 5% A; 5 min, 35% A; 7 min, 55% A; 9 min, 95% A; 10 min, 5% A; 12 min, 5% A. Violacein, with a retention time of 2.63 min and deoxyviolacein, with a retention time of 3.03 min were analysed by peak area integration at 565 nm using a standard curve.

### Flow cytometry

Flow cytometry was used to determine the composition of the cocultures at different time points. Cell fluorescence was measured using an Attune NxT flow cytometer (Thermo Scientific) using the following parameters: FSC 660V, SSC 500 V, violet laser VL1 (405 nm ex./440(50) nm em.) 420 V, blue laser BL1 (488 nm ex./530(30) nm em.) 450 V, yellow laser YL2 (561 nm ex./620(15) nm em.) 560 V. 10,000 cells were counted for each sample and the data was analysed using FlowJo. The *E. coli* population was gated using the FSC-H and SSC-H channels and singlets were identified using the FSC-H and FSC-A channels. The GFP and RFP tagged populations were separated by plotting the BL1-H and YL2-H channels. The gating used for determining the coculture composition is shown in Supplementary figure 4.

## Results and Discussion

### Identifying pathway splitting candidates

The violacein bioproduction pathway is a five-gene pathway downstream of tryptophan (Figure 1A). As tryptophan is vital for the production of violacein, we constructed a plasmid with the genes TrpED expressed under the control of a pTet promoter (pHM068), in a tryptophan-overproducing strain of BL21(DE3), referred to in this text as the B6 strain [9] (Supplementary Figure 1). We introduced the genes *vioA, vioB, vioC, vioD* and *vioE* expressed under the control of a low strength mutant T7 inducible promoter, G6 [11], on a plasmid, along with pHM068 into the B6 strain. On performing growth culture experiments, we found that there is significant growth defect (p*<*0.0001) on the expression of the pathway as compared to the wild type strain (Figure 1C). This demonstrates the genetic and metabolic burden on the cell on the introduction of the pathway. To tackle this metabolic burden, we selected three different pathway splitting points where the first strain produces an intermediate, which is transported out of the first strain and is taken up by the second strain expressing the rest of the pathway, to produce the final product, violacein. In each of these cocultures, the first strain is the Trp strain wit the TrpED sfGFP plasmid (pHM068) and the second strain is the BL21(DE3) strain with the mRFP plasmid (pHM120) and the violacein genes are expressed on a plasmid.

The three splitting strategies are Trp:VioABEDC, TrpVioABE:VioDC, TrpVioABED:VioC (Figure 1B). We used the monoculture that expressed the entire pathway in the tryptophan overproducing Trp strain as a control. The strains were grown in a microplate reader to measure their growth behaviour(Figure 1C). We saw that while the introduction of the entire violacein pathway significantly reduces the growth rate of the monoculture strain, splitting the pathway alleviates this growth defect, resulting in less pronounced growth reductions compared to the monoculture. The Trp:VioABEDC split was selected because tryptophan is a widely used metabolite in the cell and an important branching point through which metabolic flux is diverted into the violacein pathway.

Tryptophan has also been shown to traverse the cell membrane via well characterized transporters in *E. coli* [5]. The TrpVioABE:VioDC split was selected because in this split both strains were expected to have similar metabolic burden levels and hence similar growth rates. We also chose the TrpVioABED:VioC split due to the nature of the pathway, if the first cell produces PVA which is transported to the second strain, this would allow for the production of pure violacein, without deoxyviolacein, which is proven to be challenging to produce using microbial bioproduction [9].

### Investigation of different pathway split mechanisms

We then constructed the coculture strains with the violacein pathway genes expressed under the control of the inducible low strength mutant T7 promoter, G6 [11]. To monitor the composition of the coculture, the two strains were also tagged with *sfGFP* and *mRFP* respectively, expressed constitutively on a separate plasmid. The coculture strains constructed were then tested in comparison to the monoculture strain for violacein production. The three cocultures developed were inoculated with an initial ratio of 1:1 for the two subpopulation strains, cultured for 24 hours and violacein was extracted. This experiment was conducted at two different tryptophan expression levels, with and without the overexpression of TrpED. For inducing the tryptophan overproducing genes, the culture was supplemented with 100 nM aTc. We found that the TrpVioABE:VioDC split without TrpED overexpression produced the highest violacein titres (25.4 mg/L) among the cocultures and achieved slightly higher titres (not statistically significant) as compared to the monoculture (17.3 mg/L) (Figure 2A). Furthermore, we found that the TrpVioABE:VioDC had similar final cell concentrations as the monoculture, which suggests that the productivity of the violacein producing cells is higher than that of monocultures (Figure 2B). We found that the final titres of the cultures with the overexpression of TrpED were comparable to the case without induction of TrpED, but there were significant growth defects for the monoculture (p*<*0.05) on the overexpression of TrpED. This suggests that tryptophan concentration may not be the limiting factor for violacein production, as overexpressing TrpED does not lead to increased violacein titres. Hence, as the violacein titres as well as the cell concentrations were lower for the cultures with TrpED overexpression, for the following section we proceeded with the cultures without TrpED overexpression.

**Figure 2:**
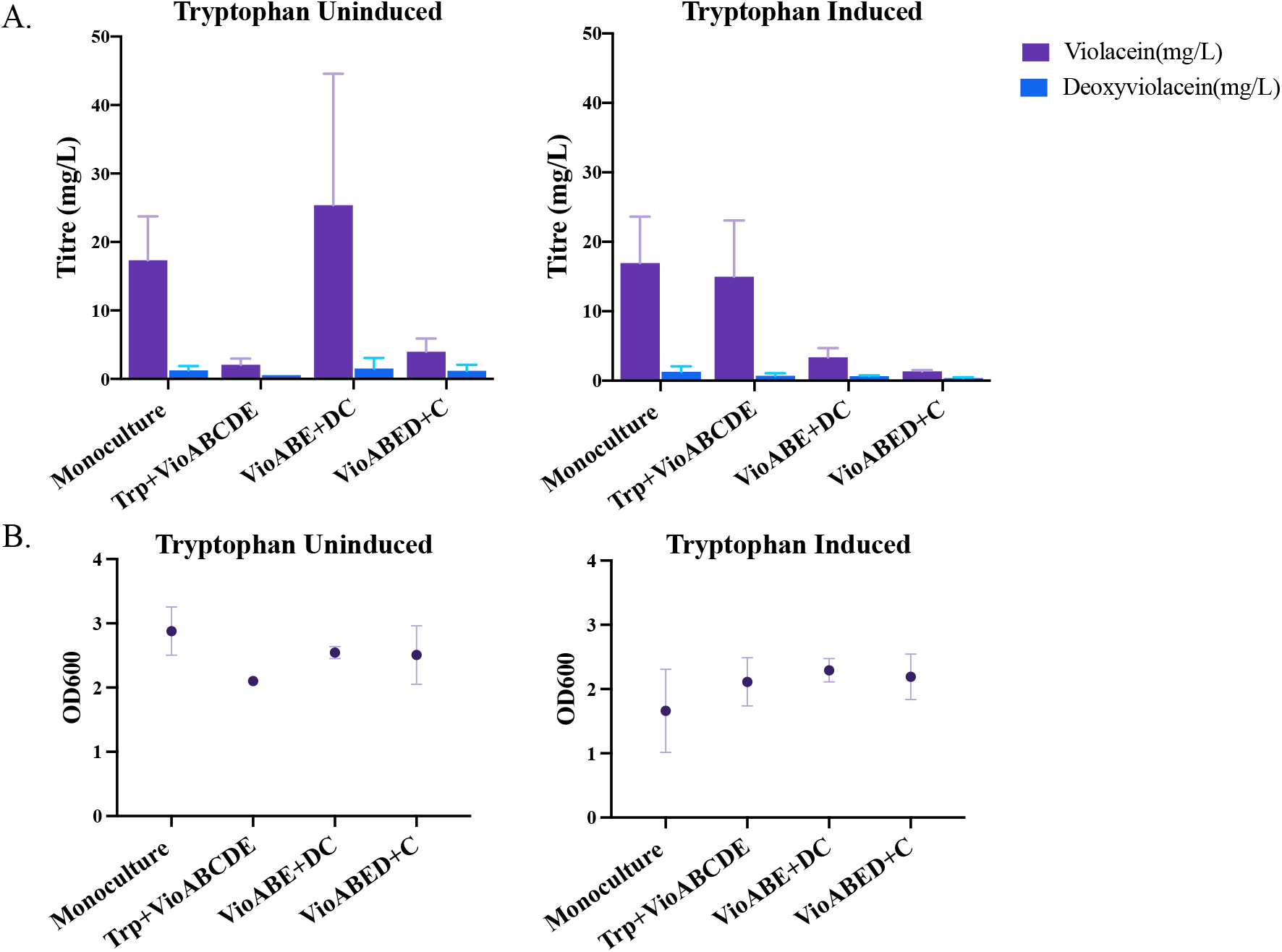
Violacein production by monoculture vs different pathway split coculture with and without overexpression of tryptophan. A. Violacein and deoxyviolacein titres for the monocultures and cocultures. B. Final cell concentrations attained by the cultures at the end of experiment at 24 hours. For tryptophan induced case, the culture was supplemented with 100 nM aTc at the time of subculture. All experiments were conducted in 5 mL liquid culture in 14 mL culture tubes. All cocultures started with 1:1 initial composition at the time of subculture. Values shown are n=3 biological replicates with mean value shown and error bars representing standard deviation. Statistically significant differences were determined using two-tailed Student’s t-test. The differences between violacein titres of the monoculture and VioABE+DC cocultures in the tryptophan uninduced condition is not significant. The differences between violacein titres of the monoculture and Trp+VioABEDC cocultures in the tryptophan induced condition is not significant.

### Coculture ratios optimize production

In order to identify the optimal coculture conditions for violacein production, we tested several coculture inoculation ratios for the three coculture systems. We tested the following initial coculture compositions for the three coculture systems: 1:9, 1:3, 1:1, 3:1 and 9:1. We found that, as with 1:1 inoculation ratio, the TrpVioABE:VioDC split performed the best of the three splits (Figure 3A). Moreover, the TrpVioABE:VioDC split coculture starting at 3:1 ratio, had the highest violacein titres (30.3 mg/L), producing significantly higher violacein titres (p*<*0.05) than the monoculture (17.3 mg/L).

**Figure 3:**
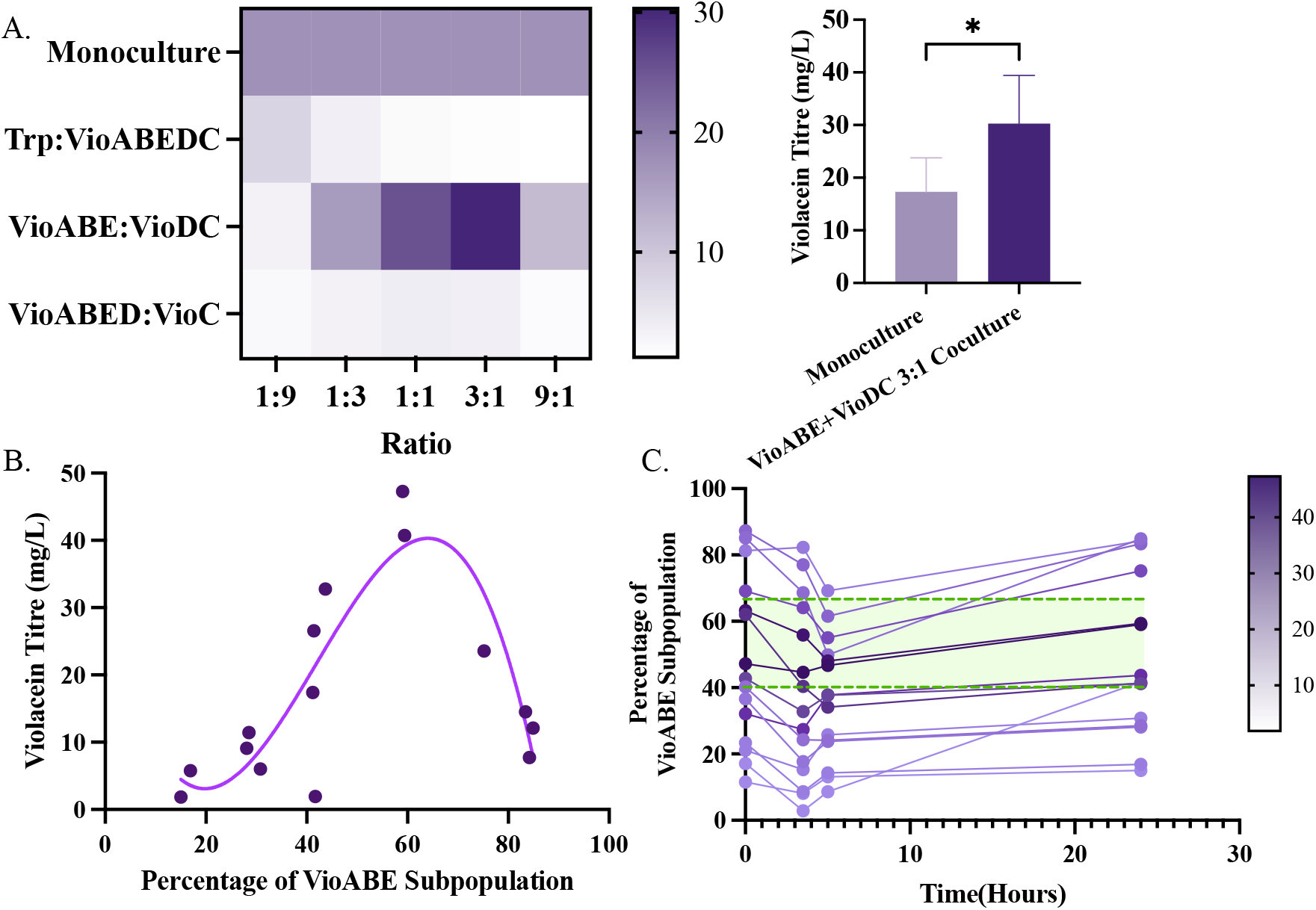
Coculture composition controls the violacein titres. A. The three cocultures Trp:VioABEDC, VioABE+VioDC and VioABED+VioC were cultured with five initial inoculation ratios of 1:9, 1:3, 1:1, 3:1 and 9:1. We see that the VioABE+VioDC split produces significantly more violacein than the monoculture. Values shown are mean violacein titres of three biological replicates. B. Violacein titres plotted against the percentage of the VioABE subpopulation in the VioABE:DC coculture. The violacein titre shows a non-liner (third order polynomial) correlation with R squared value = 0.74. C. The composition of the VioABE+VioDC coculture trajectories over the course of the experiment. The colours of the trajectories represent the violacein titres(mg/L) of the coculture. The cocultures with starting compositions between 40:60 and 65:35 produce higher titres, depicted here in green. Statistically significant differences were determined using two-tailed Student’s t-test (* represents p*<*0.1)

Based on the results for the TrpVioABE:VioDC coculture, we proceeded to identify the ideal coculture compositions for maximum violacein titres. Figure 3B shows the correlation between the final violacein titres and the final coculture composition after 24 hours of culturing. We found that the final coculture composition has a very strong impact on the final violacein titres. We observed that the final composition of 60:40 ratio produced the highest violacein titres for the TrpVioABE:VioDC coculture split. This outcome may result from a balanced conversion of intermediates and effective metabolite exchange. At lower ratios, insufficient intermediate accumulation might limit the VioDC strain’s ability to convert intermediates into violacein, while at higher ratios, the reduced number of VioDC cells could limit maximum conversion efficiency. Furthermore, we saw that the ideal coculture composition for maximum violacein production is different for different splits (Supplementary Figure 3). We also observed that the coculture composition changes through the course of the experiment (Figure 3C). We found that the cocultures diverge from the inoculation composition and very similar starting compositions can lead to different final compositions as well. For the TrpVioABE:VioDC coculture, we see that for the maximum violacein titres, the most suitable range of initial coculture composition was found to be between 40:60 to 65:35 (Figure 3C).

While the TrpVioABE:VioDC coculture split produced violacein titres comparable to the mono-culture, the other two splits Trp:VioABEDC and TrpVioABED:VioC have lower violacein titres. For the Trp:VioABEDC split, the highest violacein production is observed for the starting composition ratio of 1:9 (Figure 3A). This can be explained by the fact that as all of the violacein producing genes are expressed in the second strain, there needs to be sufficient concentrations of the second strain for maximum production. Moreover, the overproduction of tryptophan from the first strain did not compensate for the reduced concentration of the second strain in the coculture. We also found that as the growth rate of the Trp strain is higher than the growth rate of the VioABEDC strain, over time as the concentration of the Trp strain increases in the coculture (Supplementary Figure 3). In the case of the TrpVioABED:VioC coculture, for violacein production by the VioC strain, the TrpVioABED strain needs to produce sufficient PDA and it needs to be exported out of the TrpVioABED strain and imported into the VioC strain. This might be the limiting step here, leading to inefficient violacein production by the coculture. The results from the coculture experiments with the over expression of the TrpED genes can be found in Supplementary Figure 2.

## Conclusions

With the advancement in the use of synthetic microbial communities, there is a rise in the development of control systems which allow the tuning of coculture composition. There is very limited study on the need of dynamic composition control for applications in bioproduction using microbial communities. In this study, we demonstrated the effect of the coculture composition on the final product titres, and demonstrated the need for dynamic composition control for reproducible robust bioproduction of high value compounds.

In this study we systematically screened the solution space for division of labour for violacein production. Here, we examined key variables for optimizing division of labour, i.e., different pathway splitting points and coculture compositions to identify conditions where the coculture is optimised and produces the highest titres of Violacein. Among the tested splitting strategies, the pathway split TrpVioABE:VioDC emerged as the most effective, producing significantly higher violacein titres compared to other splits and the monoculture. The coculture produced the highest titres (30.3 mg/L) at a final composition around 60:40, which is significantly higher (p*<*0.05) than the titres achieved for the monoculture (17.3 mg/L). We also found that different coculture splits exhibit different trends of violacein titres at different compositions. We demonstrated that a coculture performing division of labour can be more advantageous than a monoculture when two conditions are satisfied. First, that the burden is well distributed between the two subpopulations such that the strains have similar growth rates and both the subpopulations are present at high enough concentrations, and second, that the intermediate where the pathway is split is efficiently exchanged between the two subpopulations.

## Acknowledgments

Figures were created using Biorender.com. G.-B.S. acknowledges the support received from the Royal Academy of Engineering through the RAE Chair in Emerging Technologies RAEng CiET 1819/5. J.J. acknowledges the support received from the Biology and Biotechnology Research Council (BBSRC) through the grant BB/T011289/2 as part of the ERA-CoBiotech project MIPLACE. R.L-A. received funding from BBSRC (BB/R01602X/1, BB/T013176/1, BB/T011408/1 - 19-ERACoBioTech-33 SyCoLim, BB/X01911X/1, BB/Y008510/1 – Engineering Biology Hub for Microbial Foods), EPSRC (AI-4-EB BB/W013770/1, and EEBio Programme Grant EP/Y014073/1), Yeast4Bio Cost Action 18229, European Research Council (ERC) (DEUSBIO - 949080), the Bio-based Industries Joint (PERFECOAT-101022370) under the European Union’s Horizon 2020 research and innovation programme and the European Innovation Council (EIC) under grant agreement No. 101098826 (SKINDEV). Imperial College London UKRI Impact Acceleration Account (EPSRC –EP/X52556X/1, BBSRC -BB/X511055/1). Thanks to the Bezos Earth Fund through the Bezos Centre for Sustainable Protein (BCSP/IC/001).

