## Supplementary Information for "Investigating The Potential of Division of Labour in Synthetic Bacterial Communities for the Production of Violacein"

### Supplementary Table 1 Plasmids used in this study

| Plasmid No | Plasmid Name | Description | Antibiotic |
|---|---|---|---|
| pHM068 | TrpED_sfGFP_pAB351 | TetRpTet-RBSc33-TrpED-T1+J23116-RBSc33-sfGFP-T1 in pAB351 | Amp |
| pHM053 | VioABEDC_pStA212 | T7(G6)-VioA-T7(G6)-VioB-T7(G6)-VioE-T7(G6)-VioC-T7(G6)-VioD in pStA212 | Km |
| pHM063 | VioABE_pStA212 | T7(G6)-VioA-T7(G6)-VioB-T7(G6)-VioE in pStA212 | Km |
| pHM066 | VioDC_pStA212 | T7(G6)-VioC-T7(G6)-VioD in pStA212 | Km |
| pHM064 | VioABED_pStA212 | T7(G6)-VioA-T7(G6)-VioB-T7(G6)-VioE-T7(G6)-VioD in pStA212 | Km |
| pHM065 | VioC_pStA212 | T7(G6)-VioC in pStA212 | Km |
| pHM120 | mRFP_pJBL5945 | J23116-RBSc33-mRFP-T1 in pJBL5945 | Amp |

### Supplementary Table 2 Strains used in this study

| Strain | Description | Reference | Source |
|---|---|---|---|
| DH10B | F$^-$ *mcrA* Δ(*mrr-hsdRMS-mcrBC*) φ80 *lacZ* ΔM15 Δ*lacX74 recA1 endA1 araD139* Δ(*ara-leu*)7697 *galU galK* λ$^-$ *rpsL* (Str$^r$) *nupG* | | |
| DH5α | F$^-$ φ80 *lacZΔM15* Δ(*lacZYA-argF*) *U169 recA1 endA1 hsdR17* (rK$^-$, mK$^+$) *phoA supE44* λ$^-$ *thi-1 gyrA96 relA1* | | |
| BL21(DE3) | F$^-$ *ompT hsdSB* (rB$^-$ mB$^-$) *gal dcm* (DE3) | | |
| B6 | F$^-$ *ompT hsdSB* (rB$^-$ mB$^-$) *gal dcm* (DE3) Δ*trpR* Δ*tnaA* | | (Fang et al., 2015) |
| Trp | pHM068 pStA212 TrpED_pTet_TetR + sfGFP pAB351 in B6 | HM_174 | This study |
| VioABE | pHM063 pHM068 VioABE TrpED in B6 | HM_139 | This study |
| VioABED | pHM064 pHM068 VioABED TrpED in B6 | HM_142 | This study |
| VioABEDC (Monoculture) | pHM053 pHM068 VioABECD TrpED in B6 | HM_145 | This study |
| VioC | pHM065 pHM120 VioC mRFP in BL21(DE3) | HM_262 | This study |
| VioD | pHM066 pHM120 VioCD mRFP in BL21(DE3) | HM_263 | This study |
| VioABEDC | pHM053 pHM120 VioABECD mRFP in BL21(DE3) | HM_264 | This study |

**Supplementary Table 3 Oligos used in this study**

| Oligo No | Oligo Name | Sequence |
|---|---|---|
| HM_030 | TrpED_Fwd | AAGGGGTTGGTCTCATGTGGCTCTTCGATGATGCAAACACAAAAACCGACTCTCG |
| HM_031 | TrpED_Rev | CAGTGTTGGGTCTCTGGTCGCTCTTCATTACCCTCGTGCCGCCAGTG |
| HM_032 | Ser40LeuT_Fwd | GCTGCTGGAATTCGCAGATATCGAC |
| HM_033 | Ser40LeuT_Rev | GTCGATATCTGCGAATTCCAGCAGC |
| HM_034 | Met293ThrC_Fwd | CCCAGCCCGTACACGTTTTTATGCAG |
| HM_035 | Met293ThrC_Rev | CTGCATAAAAACGTGTACGGGCTGGG |

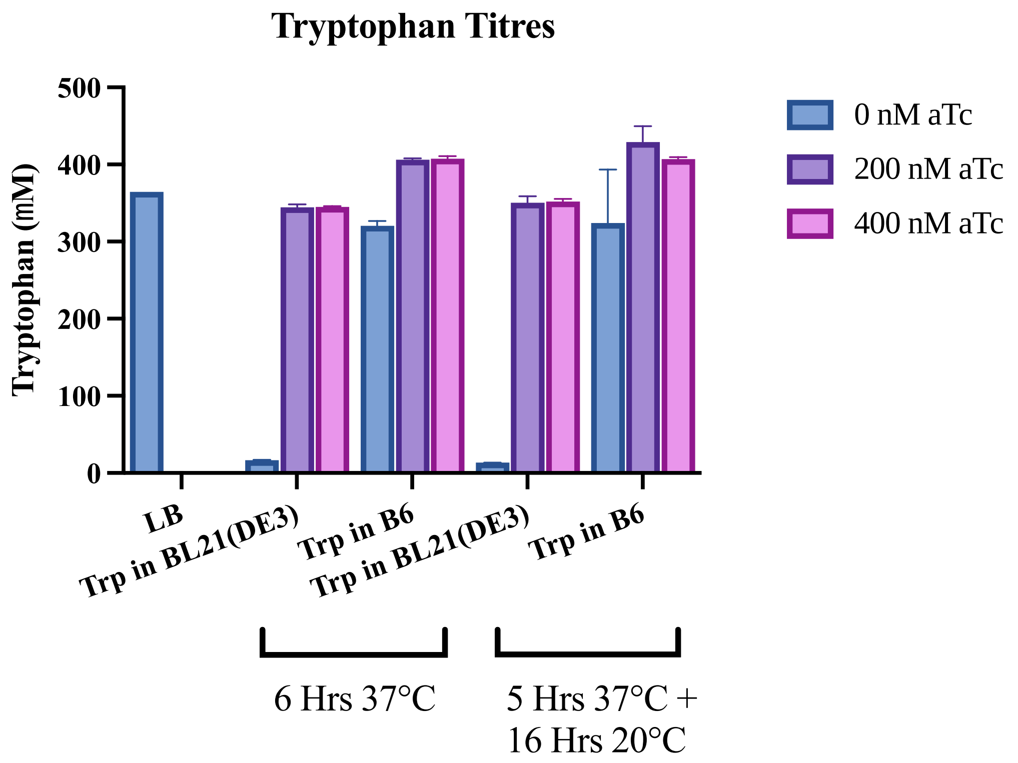


**Supplementary Figure 1**. Tryptophan production by the tryptophan overproducing strain developed. The tryptophan overproducing plasmid, pHM068, was transformed in both wild type BL21(DE3) strain and the tryptophan accumulating mutant strain B6. 2 mL of liquid cultures were incubated in 14mL culture tubes under two conditions, 6 hours at 37°C and 5 hours at 37°C followed by 16 hours at 20°C. Following this, the cells were disrupted and the supernatant was collected for tryptophan measurement by HPLC. Values shown are n=2 biological replicates with mean value shown and error bars representing standard deviation.

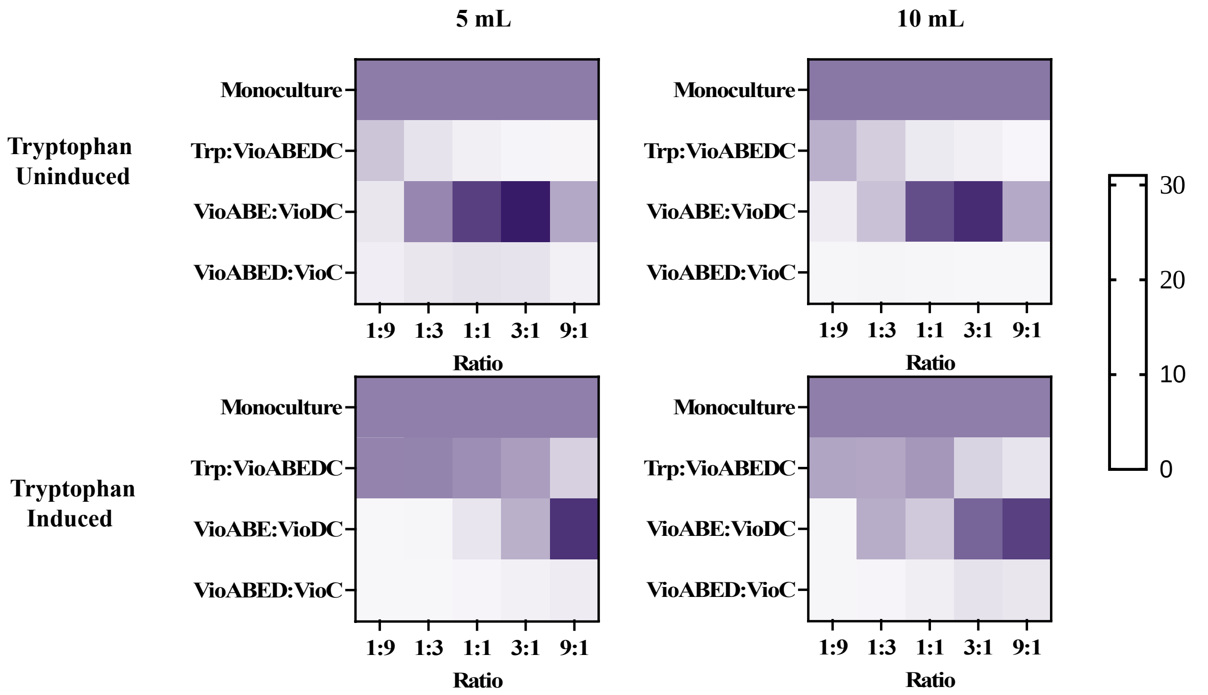


**Supplementary Figure 2**. Violacein titres produced by cocultures with different pathway split and different initial composition. The cocultures were tested at different culture volumes, 5 mL of liquid culture in 14 mL culture tubes and 10 mL of liquid culture in 50 mL shake flasks. The experiment was also repeated with and without the induction of the tryptophan overproduction genes. Values shown are mean violacein titres of three biological replicates.

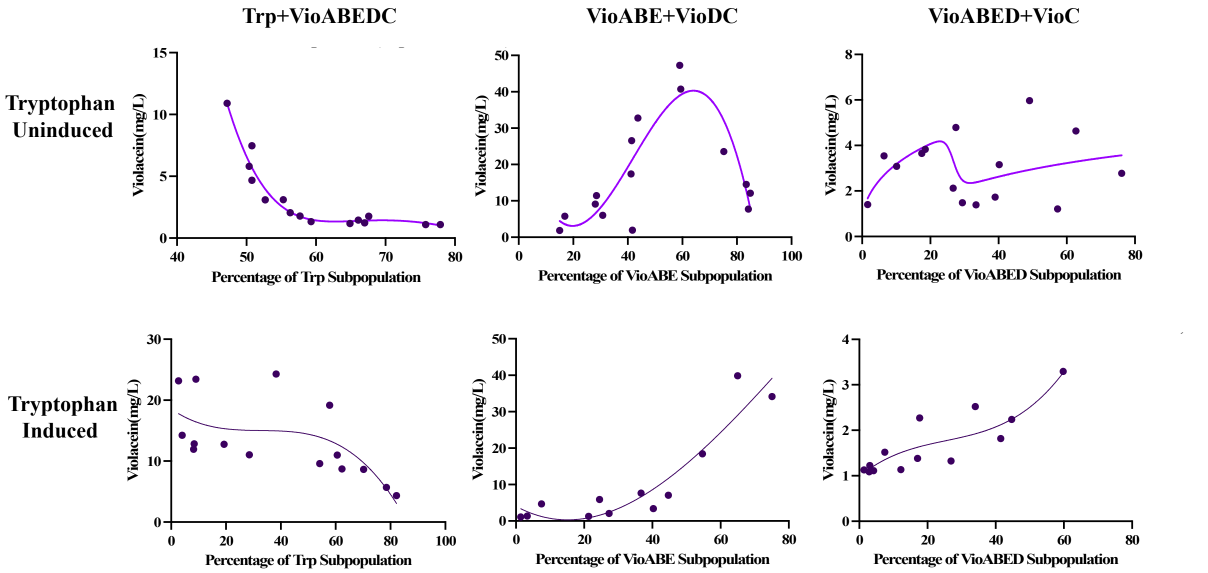


Supplementary Figure 3. Relationship between violacein tires and final coculture compositions for the different coculture splits, with and without the induction of the tryptophan overproducing genes. Lines represent nonlinear (third order polynomial) fit for the data. The R squared values for the curves in the six plots respectively are: 0.95, 0.74, ,0.46, 0.88 and 0.77.

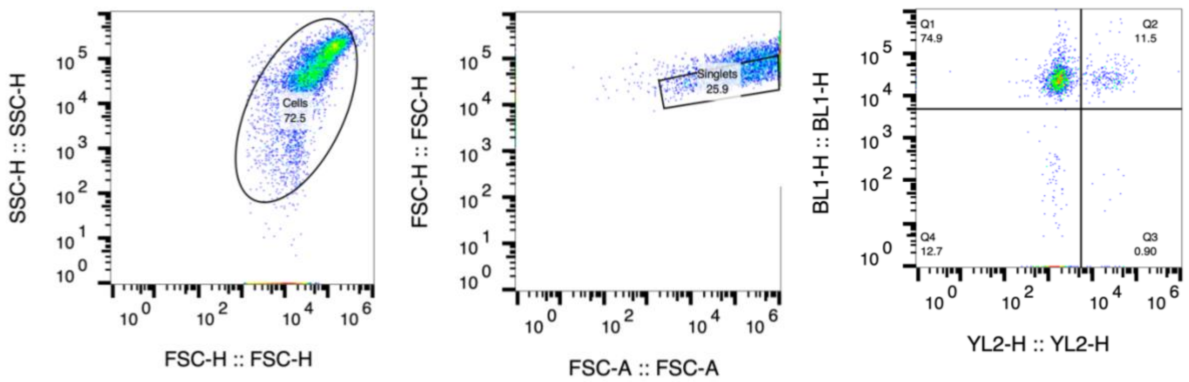


Supplementary Figure 4. Gating strategy applied for determining the coculture composition. The BL1 channel identifies the GFP tagged population while the YL2 channel identifies the RFP tagged population. The population showing positive for both RFP and GFP was identified to be cells sticking together and was hence the Q2 events were distributed equally to the GFP and RFP populations.
